# Chromatin accessibility changes at intergenic regions associates with ovarian cancer drug resistance

**DOI:** 10.1101/2021.02.24.432641

**Authors:** John Gallon, Erick Loomis, Edward Curry, Nicholas Martin, Leigh Brody, Ian Garner, Robert Brown, James M. Flanagan

**Affiliations:** Ovarian Cancer Action Research Centre, Dept Surgery & Cancer, Imperial College London, London W12 8EE, UK; Trace Element Laboratory, Charing Cross Hospital, Imperial College NHS Trust, London, UK; Desktop Genetics, 28 Hanbury St, London E1 6QR, UK; Institute of Cancer Research, Sutton, London SM2 5NG, UK

**Keywords:** cancer, chemotherapy, drug resistance, epigenomics, ovarian

## Abstract

We have investigated how genomic distribution of chromatin accessibilities alter during acquisition of resistance to carboplatin-based chemotherapy using matched ovarian cell lines from high grade serous ovarian cancer (HGSOC) patients before and after becoming clinically resistant to platinum-based chemotherapy. Resistant lines show altered chromatin accessibility at intergenic regions, but less so at gene promoters. Super-enhancers, as defined by clusters of cis-regulatory elements, at these intergenic regions show chromatin changes that are associated with altered expression of linked genes, with enrichment for genes involved in the Fanconi anemia/BRCA DNA damage response pathway. Further, genome-wide distribution of platinum adducts associates with the chromatin changes observed and distinguish sensitive from resistant lines. In the resistant line, we observe fewer adducts around gene promoters and more adducts at intergenic regions. Thus, chromatin changes at intergenic regulators of gene expression are associated with *in vivo* derived drug resistance and Pt-adduct distribution in patient-derived HGSOC drug resistance models.

## Introduction

Platinum-based chemotherapeutics, such as cisplatin and carboplatin, are clinically important first line therapies in the treatment of a wide variety of solid cancers (1, 2). These drugs exert their DNA damaging, cytotoxic effect by the formation of platinum-DNA adducts, inter- and intra-strand crosslinks, which induce cell death through a number of pathways if the adduct is not repaired (3). While many patients initially respond to platinum-based chemotherapy, they will eventually relapse with disease that fails to respond to treatment leading to poor survival (3, 4). Understanding the changes that occur in platinum resistant tumours is essential for rational approaches to circumvent resistance and developing molecularly targeted agents for use in recurrent cancers.

Epigenetic mechanisms play a key role in the development of platinum-resistance and drug tolerance (5). Cells surviving cisplatin exposure, such as transient drug tolerant persisters, or drug resistant cells surviving cisplatin selection have genome wide epigenetic alterations (5–8). Drug-tolerant persisters exhibit a repressed chromatin state and can serve as founders for further genetic and epigenetic change leading to resistance (7, 9–11). The promoters of genes susceptible to hypermethylation in ovarian tumours during the emergence of resistance to platinum resistance are marked by H3K27 and H3K4 bivalent methylation domains present in tumour cells pre-chemotherapy, further emphasizing the relationship between chromatin states and epigenetic adaptation to platinum treatment (12). Epigenome profiling of ovarian cell line models selected *in vitro* for platinum resistance has suggested a role for distal super-enhancers (SEs) and their gene targets in maintain the transcriptional program of the platinum-resistant state (13, 14). These studies highlight the multi-factorial nature of the transcriptional states driving platinum resistance, while identifying SEs that play a critical role in transcriptional regulation of platinum resistance during in vitro selection. However, it is still poorly understood whether such changes at SEs occur following *in vivo* selection following platinum-based chemotherapy of patients’ tumours as they develop clinical resistance.

We have used matched chemo-sensitive and chemo-resistant ovarian cancer cell lines isolated from high-grade serous ovarian cancer patients before and following treatment, to examine the relationship between chromatin accessibility, platinum-DNA adduct distribution, and chemotherapy resistance. To examine further the potential functional significance of any changes observed, we have correlated chromatin changes, identified using ATAC-seq, between the lines at gene promoters, CpG islands, enhancer sequences and other genomic regions with changes in gene expression.

The platinum atom of cisplatin can form covalent bonds to the N7 positions of purine bases and the density of GG dinucleotides is considered the main factor influencing distribution of platinum adducts (15). However, chromatin states could also affect the pattern of cisplatin crosslinking (16, 17). Indeed epigenetic therapies, such as histone deacetylase (HDAC) inhibitors, can increase the accessibility of chromatin to cisplatin through global chromatin decompaction, leading to enhanced cell death (18). Given the known roles for chromatin in mediating the emergence of drug resistance and since chromatin changes might influence the formation and effect of platinum adducts, we aimed to examine platinum-adduct distribution in the genome of ovarian cells and relate this to chromatin conformation and gene expression.

We have used Pt-exo-seq to map platinum adducts, genome-wide by adapting methods for exonuclease mapping of transcription factor binding sites (19–21). Pt-exo-seq is primarily based on the inhibition of exonuclease activity by the bulky platinum adduct and the subsequent removal of the adduct by cyanide treatment, followed by next generation sequencing. A schematic of the Pt-exo-seq approach is shown in Supplementary Figure 1. This complements existing methods for examining the distribution of platinum adducts genome wide such as Damage-seq (22, 23). In Damage-seq, cisplatin damaged DNA fragments are immunoprecipitated using an antibody specific to platinum-bound DNA prior to sequencing using a library preparation protocol dependent on the stalling of DNA polymerase by DNA adducts. Fragments without adducts which are non-specifically bound by the immunoprecipitation step are removed by subtractive hybridisation prior to sequencing. This method provides accurate and strand-specific mapping of DNA damage, but is dependent on immunoprecipitation and therefore heavily influenced by the sensitivity and specificity of the antibody used and the conditions for subtractive hybridisation. Pt-exo-seq avoids these potential complications while still providing accurate and strand-specific mapping at high resolution.

## RESULTS

### Chromatin conformation in matched resistant lines following platinum-based treatment

We have studied chromatin conformation by ATAC-seq in three isogenic pairs of sensitive and resistant human ovarian cell lines. PEO1 and PEO4 were both isolated from the same patient diagnosed with high grade serous ovarian cancer (HGSOC) following platinum-based chemotherapy, but before and after the clinical development of resistance (25). PEA1 and PEA2 were isolated from ascites of a patient with HGSOC prior to platinum chemotherapy and after relapse with resistant disease (25). A2780cp70 (abbreviated to CP70) is an *in vitro* derived platinum resistant derivative of the ovarian cell line A2780 (24). The characteristics and sensitivities of the cell line to cisplatin are summarized in Supplementary Table 3.

Chromatin accessibility profiles were compared between the sensitive and resistant lines in each pair, based on ATAC-seq data produced in triplicate. ATAC-seq reads were assigned to 1Kb windows tiling across the reference genome (hg19). FPKM values (Fragments Per Kilobase of DNA, per Million mapped reads of sequencing library) were calculated for each window, in each replicate. Individual windows were called differentially accessible based on a log2 fold change in coverage in the sensitive to resistant lines > ±2 and moderated t-test FDR <0.001. Substantial alterations to the landscape of chromatin accessibility were found in all 3 resistant lines compared to their sensitive counterparts, affecting 297 windows (0.009%) of the genome in the A2780 pair (Figure 1A), 8,950 windows (0.27%) of the genome in the PEA pair (Figure 1B) and 7,298 windows (0.22%) in the PEO pair (Figure 1C) out of 3,485,781 windows examined in the genome. A matrix of pair-wise correlations of chromatin accessibility across consistently differentially accessible windows (adjusting for cell line) showed the resistant PEA2 and PEO4 lines clustering together, separately from their respective sensitive PEA1 and PEO1 counterparts (Figure 1D).

**Figure 1.**
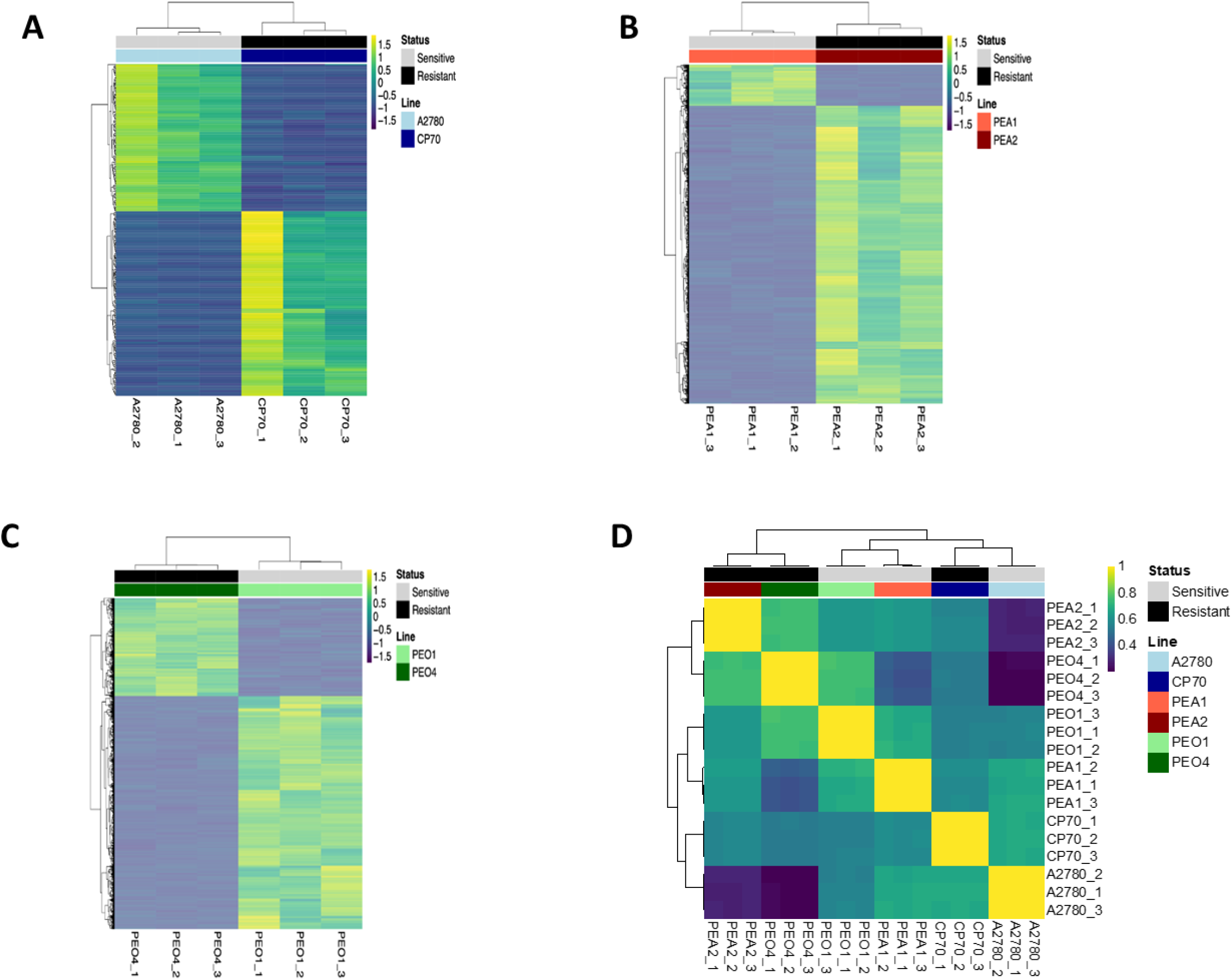
Correlation of ATAC-seq chromatin accessibility profiles. Heatmaps of ATAC-seq coverage in regions showing differential accessibility between the cisplatin sensitive and resistant line in each pair. Each row is a 1Kb window showing log2FC > ± 2 and FDR < 0.001. Dendrograms show similarity between samples as defined by unsupervised hierarchical clustering **A**) A2780/CP70, **B**) PEA1/PEA2, **C**) PEO1/PEO4. **D**. Heatmap of Spearman’s correlation coefficients for each experimental replicate for each cell line analysed, based on all windows passing filter.

Thus, the resistant HGSOC tumour cells isolated at time of recurrence with non-responsive disease show similarity, suggesting common chromatin changes occurring during the acquisition of resistance *in vivo*. The HGSOC lines were in turn different in their chromatin accessibility to the non-HGSOC, *in vitro* resistance derived, A2780-CP70 pair, potentially reflecting their different origin and *in vitro* method of drug selection.

### Platinum resistance associates with differential chromatin accessibility in intergenic regions

Each of the windows showing differential accessibility between sensitive and resistant lines by ATAC-seq was annotated by genomic class using HOMER (35). Those that were associated with genes were classed as either CpG island, promoter-TSS (−1 Kb to 100 bp 3’ of Transcription Start Site, TSS), exon, intron, 5’ untranslated region (UTR), 3’ UTR, or TTS (−100 bp to 1 Kb 3’ of TTS). The number of windows falling within each genomic class are shown in Supplementary Table 4. Those falling outside these classes were classed as intergenic. We then calculated odds ratios (OR) of enrichment for each class of genomic element in windows showing either increased or decreased accessibility in the resistant line compared with all windows (shown as log(OR) in Figure 2).

**Figure 2.**
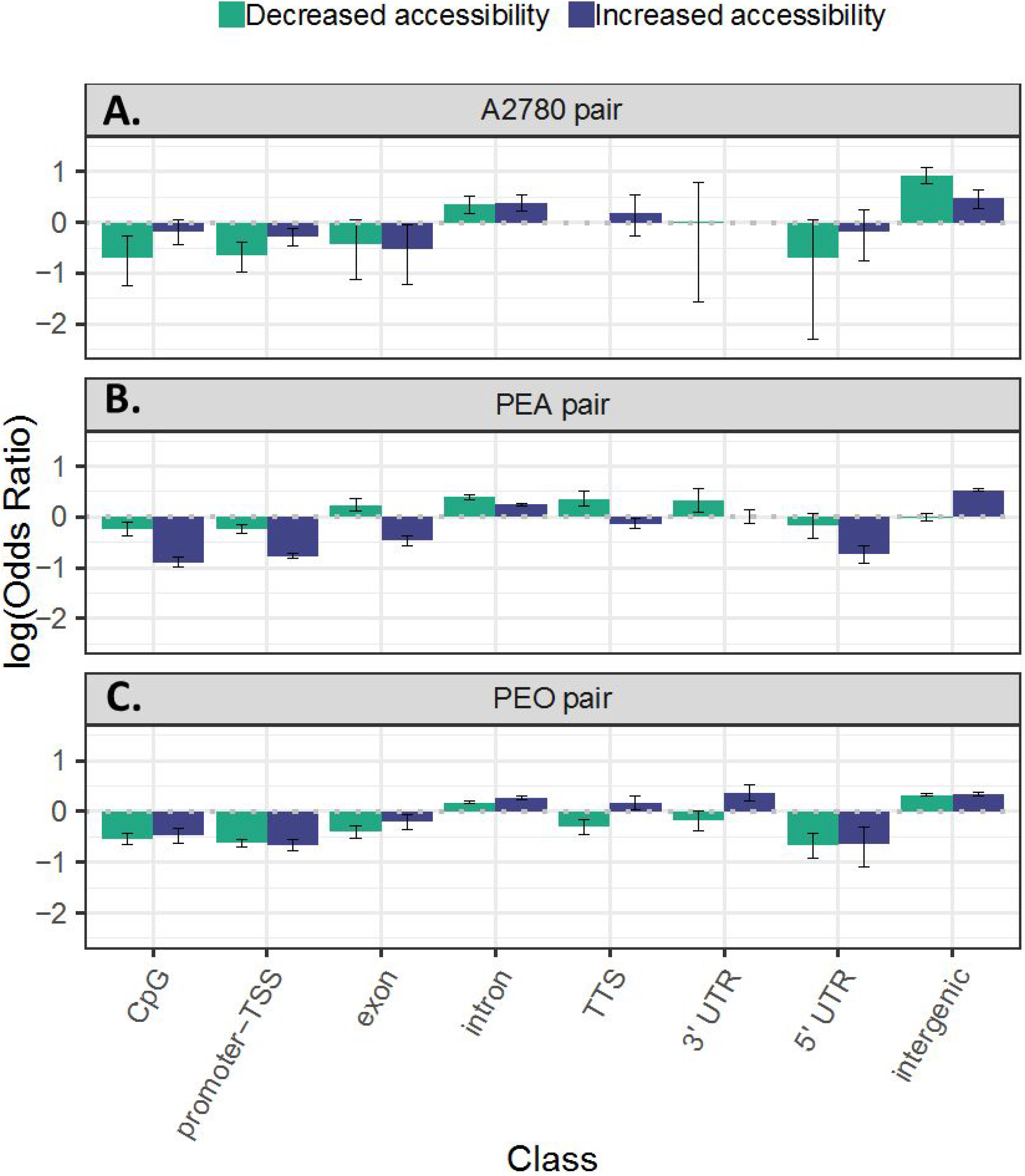
Enrichment for genomic elements in regions showing differential accessibility. Log(OR) from Fisher’s exact test for enrichment for each class of genomic element in sets of windows showing increasing or decreasing chromatin, between the sensitive and resistant lines in each pair. **A**) A2780/CP70, **B**) PEA1/PEA2, **C**) PEO1/PEO4. OR >1 or <1 corresponds to enrichment for a class. Error bars represent 95% confidence interval

In all three sensitive/resistant pairs, there was under-representation of promoter-TSS, CpG island and exonic windows in the sets of differentially accessible windows, suggesting fewer chromatin changes at these genomic regions during the acquisition of resistance (Figure 2). In all cases there was enrichment for intronic windows in the sets of differentially accessible windows, demonstrating chromatin changes at intronic regions as strongly associated with resistance. Intergenic elements were also highly over-represented in differentially accessible windows in all three pairs.

### Intersection of expression and promoter chromatin accessibility

Analysis of differential gene expression between sensitive and resistant lines was performed by RNA-seq to detect alterations to gene expression in the resistant lines. Genes were defined as differentially expressed based on a log2FC in normalized read count of > ±2 and moderated t-test FDR < 0.05. The summary of differentially expressed (DE) genes and their direction of change for each pair of cell lines is shown in Supplementary Table 5. Although many genes show changes in expression, few show universal differential expression in the same direction in all three pairs. By overlapping lists of genes showing significantly altered expression in all three resistant lines (DE = log2FC > |2|, FDR < 0.05), we identified only 4 genes which were consistently upregulated, *PARP9*, *SPHK1*, *DDX60L* and *BCAM*, while only 5 genes were consistently downregulated *CCDC80*, *TLE4*, *THBS1*, *MAP1A* and *VEPH1*. Using alternative methods of analysis (a sensitive-resistant linear model incorporating all three pairs, and a rank product analysis) only *THBS1* was consistently defined as downregulated using all methods. We confirmed the differential expression of THBS1 and PARP9 using RT-QPCR (p < 0.05, t test, Supplementary Figure 2). These results suggest there is minimal overlap between alterations of individual gene expression across these three models, although we cannot exclude more subtle changes in gene expression or gene networks being in common.

In order to relate gene expression with chromatin accessibility, ATAC-seq coverage was measured in the 2 Kb region spanning the transcription start site (TSS) of the genes defined as differentially expressed using rank product analysis in each cell line pair (Supplementary Figure 3). Genes which showed reduced expression in the resistant lines had slightly reduced accessibility compared to the resistant lines, although this did not reach statistical significance (t-test p > 0.05) (Figure 3). Genes which were upregulated in the resistant lines showed the inverse, whereby there was a stronger ATAC-seq signal around the TSS of these genes in the resistant lines in which they were upregulated, compared to the sensitive lines (t-test p < 0.01). ATAC-seq data at example genes upregulated or downregulated are shown in Supplementary Figure 3.

**Figure 3.**
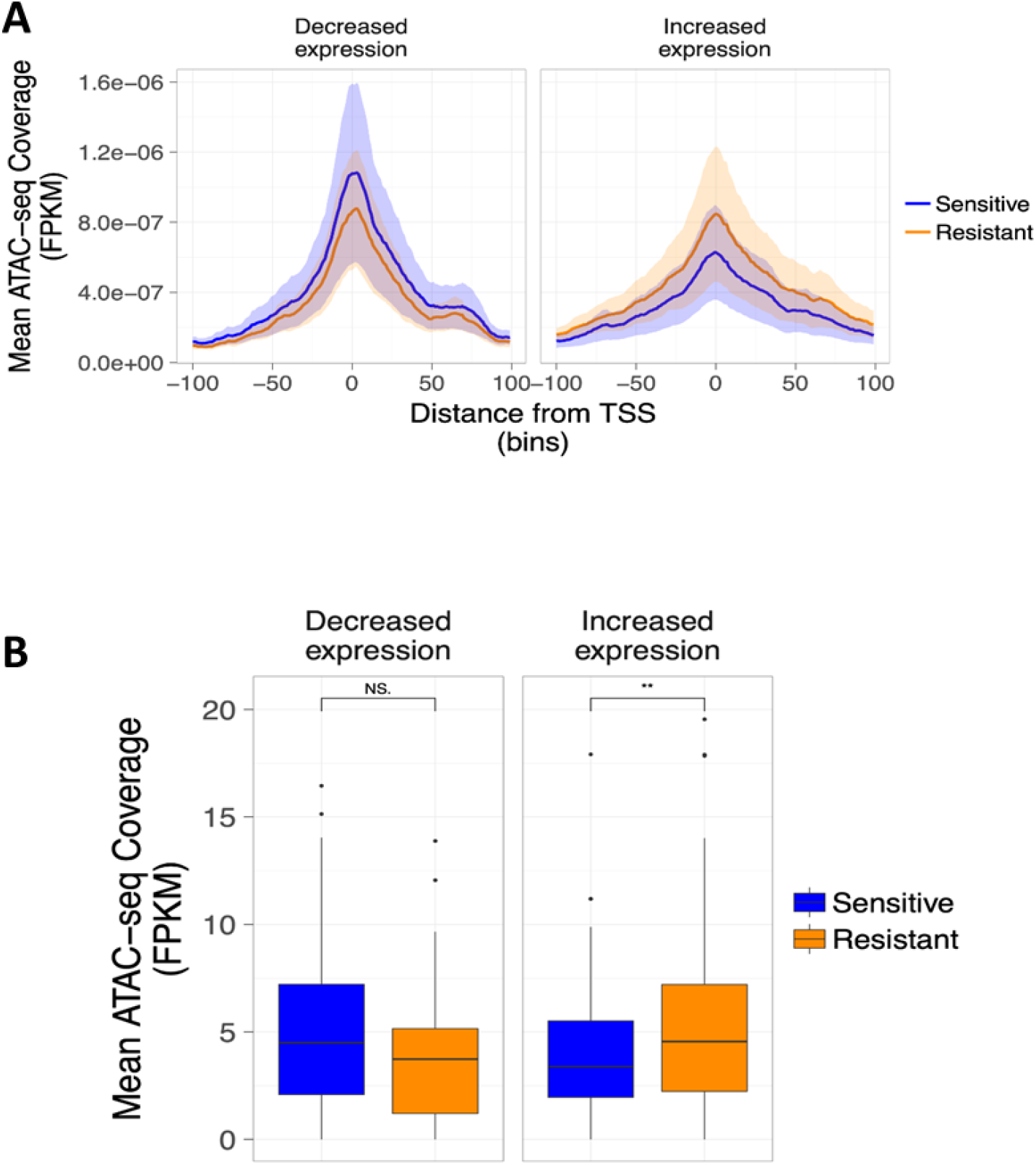
ATAC-seq coverage around TSS of differentially expressed genes. A) ATAC-seq coverage around TSS of differentially expressed genes for genes with decreased expression and those with increased expression defined by rank product analysis. B) Mean ATAC-seq coverage in 2Kb around TSS of differentially expressed genes (t test, NS, not significant; ** p< 0.01).

Thus, these trends are consistent with an accessible promoter being associated with a permissive state for gene transcription and an inaccessible promoter with transcriptional silencing. However, relatively few changes in chromatin accessibility were observed in the resistant lines at promoter regions (Figure 2) and those that did occur only weakly correlated with changes in gene expression (Figure 3).

### Chromatin accessibility change at enhancer elements associates with gene expression in resistant lines

The differences detected in chromatin conformation between sensitive and resistant lines frequently occurred at intergenic regions. Given the minimal association between chromatin accessibility changes at gene promoters and altered gene expression, we asked how changes to chromatin conformation at intergenic regions associated with transcriptional changes. The algorithm CREAM (Clustering of genomic REgions Analysis Method) was used to define clusters of cis-regulatory elements (COREs) (also known as super-enhancers) in each of the lines, based on peaks called by MACS2 from the merged alignment file produced from the 3 ATAC-seq replicates from each cell line (https://www.biorxiv.org/content/10.1101/222562v1). The results of this are summarised in Supplementary Table 6, along with the numbers of COREs unique to each line. We identified substantial changes to super-enhancer landscapes in all three resistant lines, compared to the sensitive. In both the A2780 and PEO pairs there was a loss of COREs (1030 to 358 and 715 to 537 respectively), while a gain in COREs was found in PEA2 compared to PEA1 (744 to 811), reflecting the trends in the direction of altered accessibility in each pair. Illustrative examples of super-enhancers with differential accessibility (representative ATAC-seq mapping tracks) for all three cell line are shown in Supplementary Figure 4.

Having defined COREs that were gained or lost in the sensitive/resistant pairs we examined if this change in chromatin conformation at super-enhancer regions was associated with changes in expression of linked genes. For each super-enhancer gained or lost, genes within 100 Kb were extracted from the Ensembl database and those that were differentially expressed between the sensitive and resistant lines identified from the RNA-seq data. The distribution of t-statistics was then plotted, for the comparison of expression values between RNA-seq replicates from the sensitive and resistant line in each pair. In all three pairs, loss of super-enhancers in the resistant line was associated with a significant shift towards reduced expression of genes proximal to the enhancers. When chromatin accessibility changes led to the gain of a super-enhancer in the resistant line, there was a shift towards increased expression of these genes in the resistant line compared to the sensitive (Figure 4A). We further tested the association between loss or gain of enhancers by testing for enrichment of differentially expressed genes in the 100 Kb surrounding lost/gained COREs. Figure 4B shows the odds ratios of finding differentially expressed genes near super-enhancers which were gained or lost in the resistant line compared to the rest of the genome. In the PEA pair, there was enrichment for differentially expressed genes near COREs lost (p < 0.05, OR 1.3, CI 1.04 - 1.6, Fisher’s exact test) and a strong enrichment for differentially expressed genes near COREs gained (p < 0.001, OR 1.62, CI 1.32 - 1.97, Fisher’s exact test). In contrast, there was significant enrichment for differentially expressed genes near COREs lost in PEO4 (p < 0.001, OR 1.98, CI 1.48 – 2.62, Fisher’s exact test) and to those gained (p < 0.001, OR 1.88, CI 1.31 – 2.65. Fisher’s exact test). While there was no significant enrichment for differentially expressed genes near COREs which were gained in CP70 from A2780, similar to PEO4, there was a 2.9-fold enrichment for differentially expressed genes near COREs lost in CP70 from A2780 (p < 0.001, OR 2.87, CI 2.28-3.58, Fisher’s exact test).

**Figure 4.**
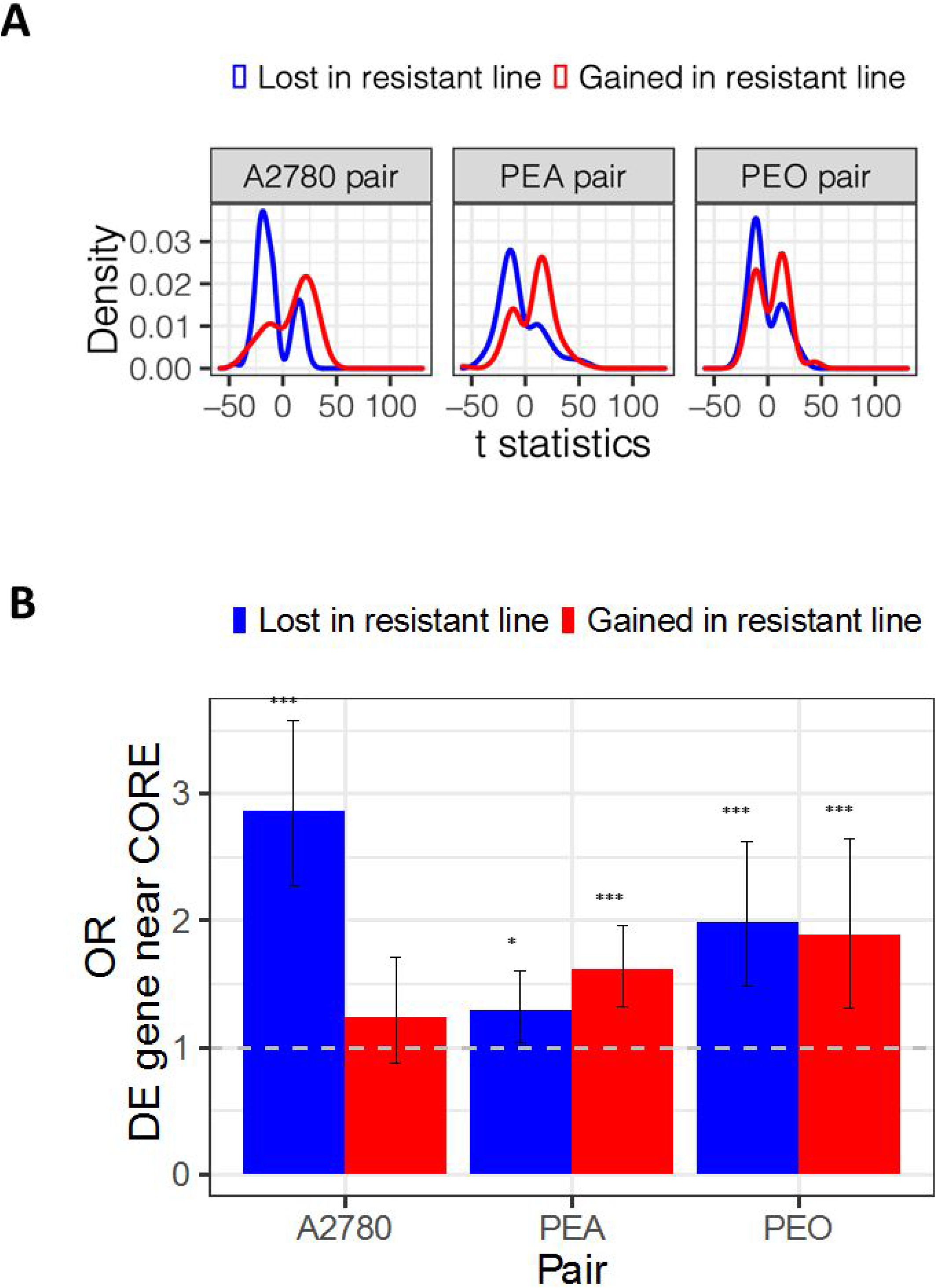
Relationship between chromatin changes at super-enhancers and gene expression. **A.** Distribution of t statistics of difference in expression for differentially expressed genes near COREs lost or gained in the resistant line in each pair. Differentially expressed genes show log2FC > ± 2 and FDR < 0.05. Red lines show distribution of t statistics for differentially expressed genes near COREs gained in resistant lines, blue lines show t statistics for differentially expressed genes near COREs lost in resistant lines. **B.** Odds ratios of differentially expressed genes being found near COREs lost or gained in resistant lines compared to their sensitive counterpart. Bars show odds ratios and error bars confidence intervals from Fisher’s exact test. * p < 0.05, *** P < 0.001.

Thus, the resistant line in all three pairs show enrichment of differential expression of genes near super-enhancers with altered chromatin accessibility. Importantly, within the set of genes near super-enhancers gained in PEO4, we observe enrichment for genes involved in the Fanconi anemia/BRCA DNA damage response (DDR) pathway (36) that are associated with clinical response to DNA damaging cytotoxics in breast cancer (p< 0.05, OR 4.8, CI 1.02 – 16.0, Fisher’s exact test), highlighting a potential mechanism through which altered super-enchancer landscapes may drive the acquisition of resistance to DNA damaging agents. Furthermore *SOX9,* recently identified as maintaining chemoresistance in ovarian cancer cells through altered accessibility at its associated enhancer (14), was among the genes near a super-enhancer which was gained in the resistant PEO4 line (see Supplementary Figure 5A).

We examined in more detail the expression of the genes included in the BIOCARTA_ATRBRCA curated list of genes involved in BRCA related cancer susceptibility. As shown in Figure 5, expression of the 19/22 genes for which we had RNA-seq data was sufficient to separate PEO1 from PEO4 and PEA1 from PEA2 by unsupervised hierarchical clustering. In the PEO pair, 12 of these genes showed small (log2FC < |1|) but significant (FDR < 0.05) changes in expression in PEO4 compared to PEO1. These included RAD51, ATM, FANCA, HUS1, BRCA1 and CHEK1 which were downregulated in PEO4. In the PEA pair, 14 of these showed a similar change in expression, of which NBN, TREX1, ATM, CHEK1, FANCC, FANCG, BRCA1 and BRCA2 were downregulated (FDR < 0.05).

**Figure 5.**
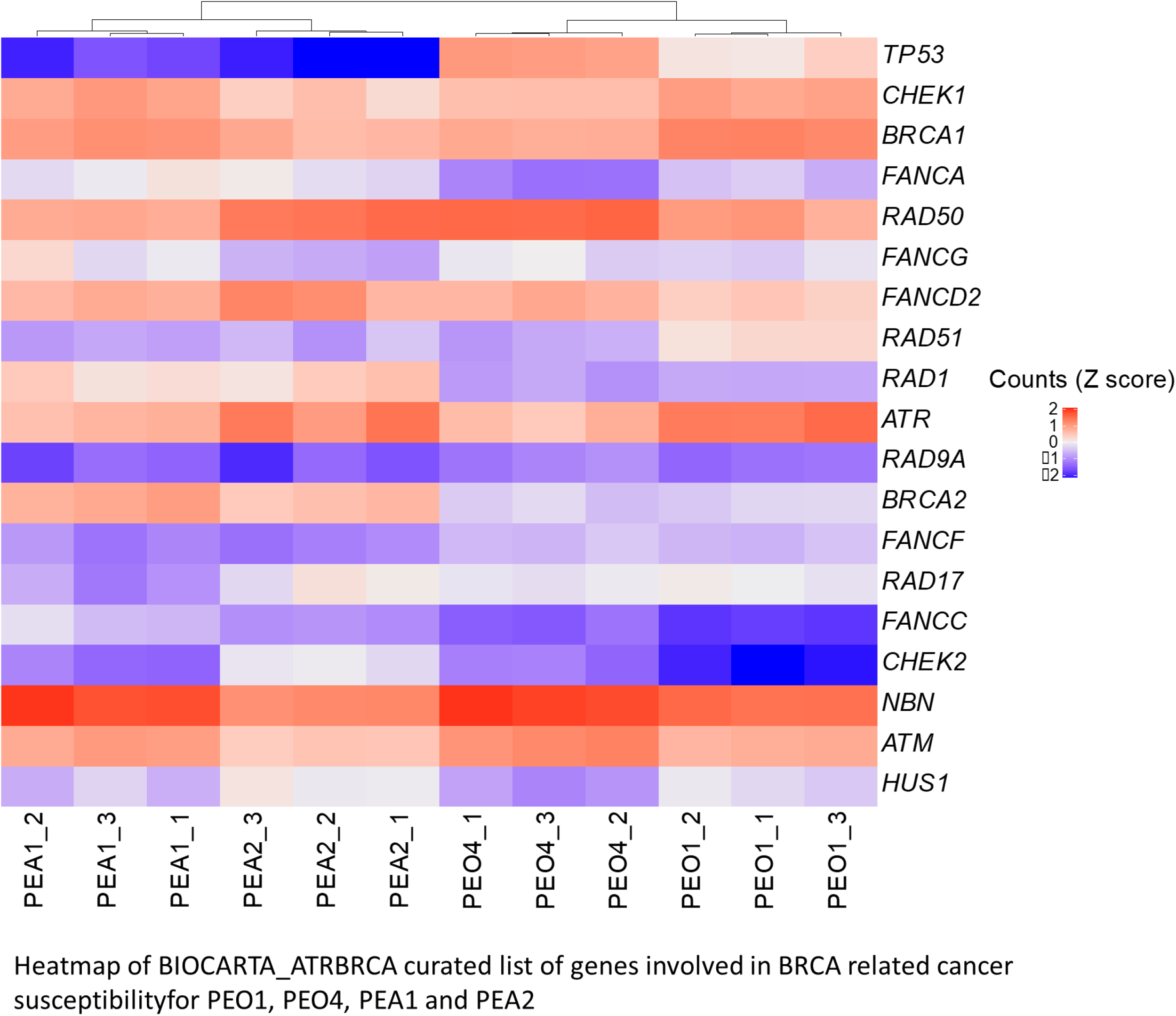
Expression of HR genes in platinum sensitive and resistant cell lines. Heatmap of Z scaled normalised counts for 19 genes in the BIOCARTA_ATRBRCA gene set, from RNA-seq from three replicates of PEO1, PEO4, PEA1 and PEA2. Dendrogram shows hierarchical clustering using the ward.D2 algorithm, based on the expression of these genes.

### Genome mapping of Pt-adducts in sensitive and resistant lines

To analyse sites of platinum adducts at high resolution, we have adapted methods for exonuclease mapping of transcription factor binding sites and developed Pt-exo-seq (20, 21). A schematic of the Pt-exo-seq approach is shown in Supplementary Figure 1. Massively parallel DNA sequencing is used to identify positions where 5’ to 3’ exonuclease digestion is blocked by platinum-DNA adducts. Levels of platinum uptake and DNA adducts formed can vary between cell lines, with many platinum resistant cell lines showing lower levels of drug uptake than their sensitive counterparts (37). While the Pt-sensitive PEA1 and Pt-resistant PEA2 lines show similar levels of Pt-adducts, as measured by Inductively coupled plasma mass spectrometry (ICP-MS), when cells are treated over a range of platinum doses, the PEO4 and A2780/cp70 resistant cell line shows markedly fewer total platinum adducts than the sensitive PEO1 and A2780 cell lines (Figure 6A). Therefore, in order to compare distribution of Pt-adducts across the genomes of PEO1 and PEO4 we have used doses of cisplatin treatment that induce approximately equivalent levels of total Pt-adducts: 16μM and 32μM respectively.

**Figure 6.**
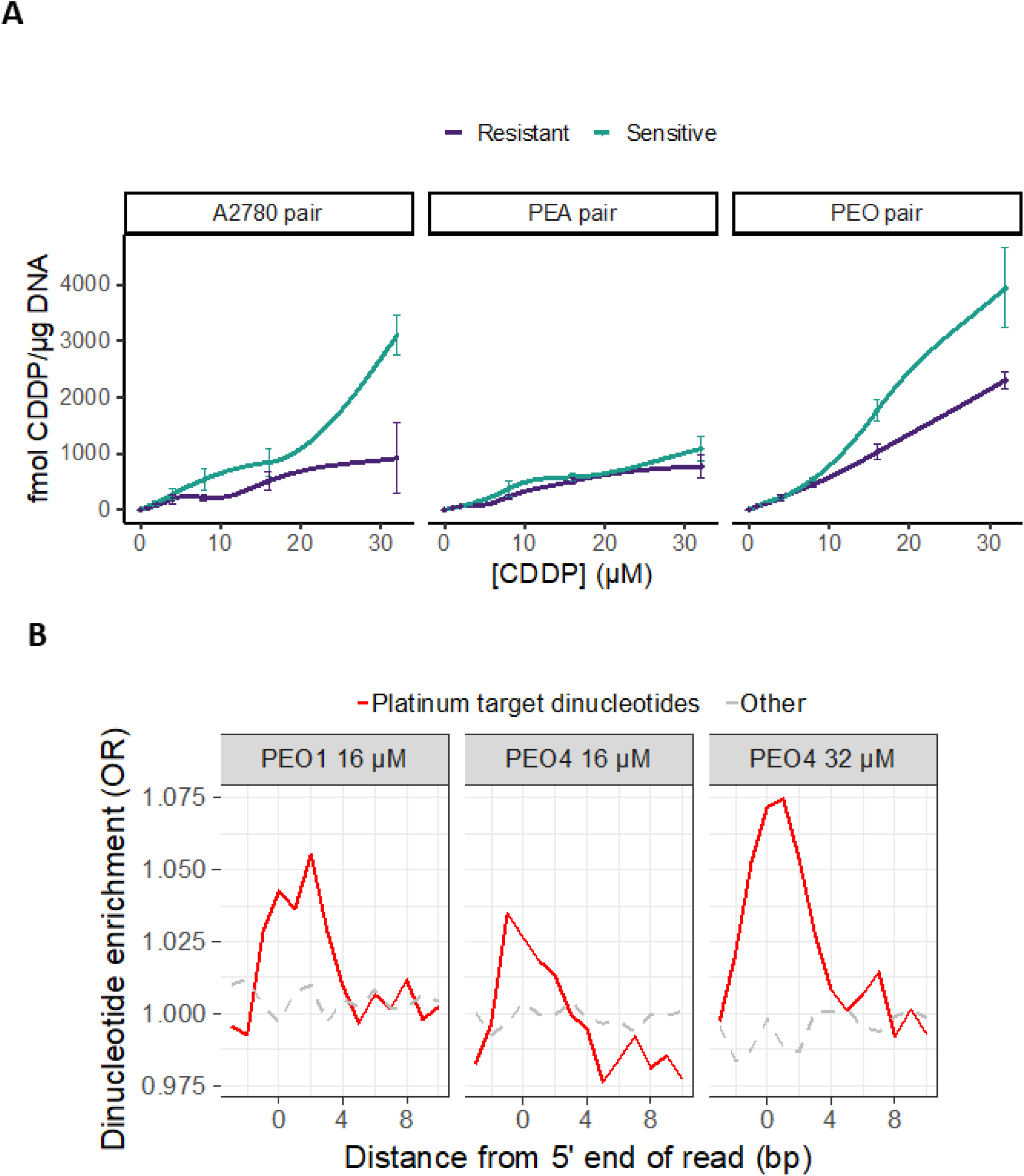
Platinum adducts in PEO1 and PEO4. (A) PEO1 incurs more adducts than PEO4 at the same dose of cisplatin, as measured by ICP-MS. (B) The difference in adduct formation is reflected in levels of enrichment for platinum target purine dinucleotides at the 5’ end of Pt-exo-seq reads. PEO1 and PEO4 were treated with concentrations of cisplatin that ICP-MS indicated would induce equal amounts of damage.

Cisplatin forms covalent bonds to the N7 positions of purine bases leading to DNA crosslinks. The majority of intrastrand crosslinks are 1,2-d(GpG) crosslinks, followed by a small number of 1,2-d(ApG) crosslinks (38). We would predict therefore that the majority of fragments detected by Pt-exo-seq will have enrichment for platinum target purine dinucleotides at the 5’ end of Pt-exo-seq reads. Indeed at doses of 100 μM cisplatin in A2780 we observe a dose dependent increase in the exonuclease resistant fraction of DNA and a 2.5 fold increase in purine dinucleotides at the terminal position (Supplementary Figure 6), which compares favourably to the 2-fold enrichment in dinucleotides previously observed using the Damage-seq assay at 200μM cisplatin. (22, 23). As shown in Figure 6B, at less highly toxic doses of cisplatin in the HGSOC lines, the 5’ end of Pt-exo-seq reads from DNA isolated from PEO1 and PEO4 cells treated with cisplatin are also enriched for purine dinucleotides compared to reads from untreated cells. At doses of 16μM cisplatin, there was less enrichment for purine dinucleotides detected following Pt-exo-seq in the resistant PEO4 line compared to the sensitive PEO1 line, while at 32μM cisplatin treatment of PEO4 there were approximately equivalent levels of enrichment for purine dinucleotides. Together these data support Pt-exo-seq as being able to detect the location of platinum adducts in genomic DNA.

### Distribution of Pt adducts in sensitive and resistant lines

Genome-wide Pt-exo-seq-coverage was calculated in 1Kb windows, as a log2FC over the untreated signal. This window size falls in the same order of magnitude as the median size of MACS2 called ATAC-seq peaks and provides sufficient depth for the signal/noise ratio to be calculated. These profiles were used to compare platinum-DNA adduct formation in PEO1 and PEO4 cells, at equal doses (16μM vs 16μM), and doses inducing equal amounts of damage (16μM vs 32μM respectively). Mean Pt-exo-seq signal was calculated for each window, across three replicates for each line and treatment condition, and used to calculate a log2FC in damage, after taking account of the untreated controls, between PEO1 and the 32μM treated PEO4 cells. We found 1,170 1Kb windows showing a log2FC > ±2 and moderated t-test FDR < 0.05. These were then separated into two groups based on direction of change in damage in PEO4 compared to PEO1; increased or decreased frequency of DNA adducts. Hierarchical clustering was then performed on the normalized read counts, using these windows, which separated the PEO1 16μM replicates from the PEO4 replicates treated at 16μM and 32μM, with the PEO4 replicates clustering by dose (Figure 7A). The PEO1 and PEO4 also separated from each other when clustering on all windows analysed (Supplementary Figure 7).

**Figure 7.**
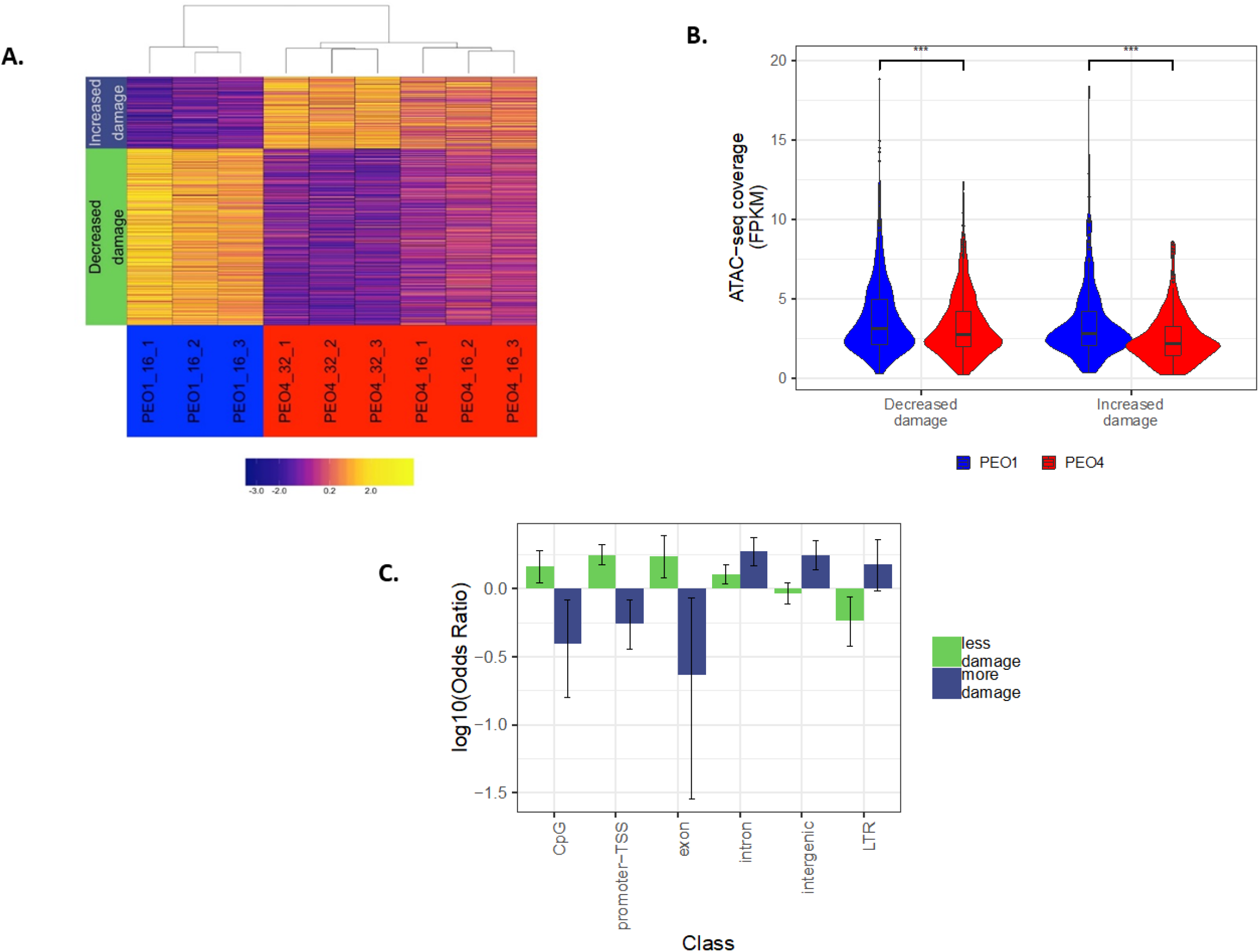
Platinum adduct distribution. (A) Heatmap of Pt-exo-seq coverage in differentially damaged windows, for PEO1 and PEO4. 1,170 1Kb windows showing mean log2FC in Pt-exo-seq signal > ±2 and FDR < 0.05, between 16 μM cisplatin treated PEO1 replicates and 32 μM treated PEO4 replicates (5 hour treatment). Dendrogram shows result of unsupervised hierarchical clustering of samples by Spearman’s correlation coefficients. Blocks on left show separation of windows into those showing either increased or decreased damaged in PEO4. (B). Chromatin accessibility in windows showing more or less platinum adduct formation in PEO4 compared to PEO1. PEO1 cells were treated with 16 μM cisplatin and PEO4 treated with 32 μM for. 5 hours. Difference in means: Decreased damage = 0.5, Increased damage = 0.90 (*** p < 0.001, Wilcoxon test). (C) Enrichment for classes of genomic elements in windows showing differential damage between PEO1 cells treated with 16 μM cisplatin and PEO4 treated with 32 μM for 5 hours. Odds ratios for only genomic classes showing significant enrichment (p<0.05) in sets of windows showing increased or decreased damage in PEO4 compare to PEO1. Error bars show 95 % confidence interval (Fisher’s exact test).

Thus, even though cisplatin is a relatively non-specific DNA damaging agent and might be expected to induce adducts at purine dinucleotides throughout the genome, there are clear and consistent differences in distribution of Pt-adducts between these matched sensitive and resistant ovarian cell lines and this is independent of the overall level of adducts formed.

### Effect of genomic context on Pt-adduct distribution

Chromatin accessibility was investigated in relation to cisplatin-DNA adduct formation. ATAC-seq coverage was calculated for each of the 1 Kb windows previously defined by Pt-exo-seq as having different levels of Pt-DNA adducts between PEO1 16μM and PEO4 32μM replicates. Thus, we aimed to assess changes in chromatin accessibility between PEO1 and PEO4 at regions that showed differential DNA Pt-adduct levels. Regions incurring fewer Pt-adducts in PEO4 showed a significant reduction in accessibility as measured by ATAC-seq in PEO4 compared to PEO1 (p < 0.001, t test) (Figure 7B). Surprisingly, when carrying out the same analysis for the smaller set of windows showing increased damage in PEO4, an even greater reduction in accessibility was observed (p < 0.001, t test).

Odds ratios were then calculated for enrichment of each class of genomic element in the sets of regions showing increased or decreased damage in the 32μM treated PEO4 samples compared to PEO1 (Figure 7C). Significant underrepresentation of regions annotated as CpG island, promoter-TSS and exon was detected in the set of regions showing increased damage in PEO4 (CpG OR 0.40, CI 0.16 – 0.83, promoter-TSS OR 0.56, CI 0.36 – 0.83, exon OR 0.24, CI 0.03 – 0.86, Fisher’s exact test). Conversely, regions showing decreased damage in PEO4 were over-represented for these same annotations. Intergenic annotated regions were enriched in the set showing increased damage in PEO4 (OR 1.80, CI 1.41 – 2.29). The decreased damage around promoters in the resistant line and the increased damage at intergenic regions suggest differential adduct formation in resistant cells is related to different genomic contexts which influence the rate of occurrence, or repair of adducts, or have implications on how they are tolerated by the cell.

To examine this further, we compared Pt-exo-seq data at COREs shared between PEO1 and PEO4, those only in PEO1 (i.e. lost in PEO4) and those only in PEO4 (i.e. gained in PEO4) (Supplementary Figure 8). Significantly more platinum adducts were detected in the COREs in the resistant PEO4 line compared to COREs in the sensitive PEO1 line. Interestingly, the COREs lost in PEO4 incurred more adducts than COREs that are maintained in PEO4 (i.e. the shared COREs). This is consistent with a hypothesis whereby DNA damage at chromatin in less accessible super-enhancers are better tolerated and could be acting as DNA damage ‘sinks’ leading to drug resistance.

### Effect of modulating chromatin accessibility on cisplatin-induced DNA adduct levels

Treating ovarian tumour cells with the HDAC inhibitor Vorinostat causes a global increase in chromatin accessibility (Figure 8A). 20μM Vorinostat treatment for 24h before chromatin extraction caused a significant increase in fragments of a length associated with mono-, di- and tri-nucleosomes following MNase digestion, consistent with a more open chromatin conformation (mono-, p < 0.01, di-, p < 0.01, tri- p < 0.05, larger, p < 0.01. t test). The same Vorinostat treatment followed by cisplatin treatment of cells, leads to a higher level of Pt-induced adducts, as measured by ICP-MS, compared to vehicle pretreatment (p < 0.001, t test) (Figure 8B). These data are consistent with a more open chromatin conformation increasing platinum DNA adduct formation. While Vorinostat treatment enhances the sensitivity of cells to cisplatin, this enhanced platinum sensitivity is observed in both platinum sensitive PEO1 and resistant PEO4 HGSOC cells (Figure 8C), suggesting a lack of specificity to drug resistant cells.

**Figure 8.**
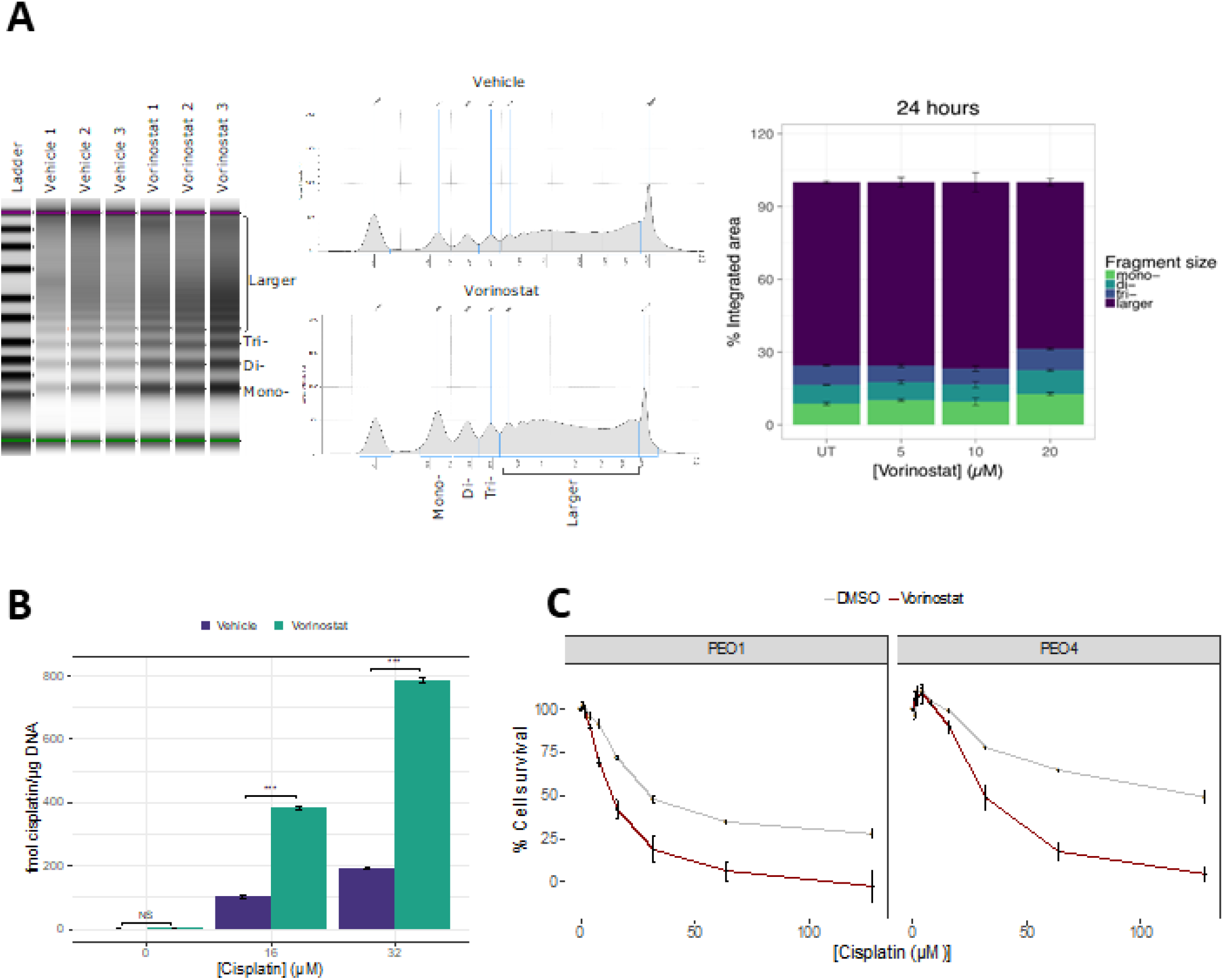
Effect of Vorinostat treatment on global chromatin accessibility and Pt-adduct formation. A) PEO4 HGSOC cells were treated with either vehicle or Vorinostat for 24h. The nucleosome distribution and percent integrated area associated with mono-, di- and tri-nucleosome DNA fragments following MNase digestion are shown. There was a significant difference in integrated area between the vehicle and Vorinostat treated cells. Mono-, p < 0.01, di-, p < 0.01, tri-p < 0.05, larger, p < 0.01). B) Levels of DNA platination as measured by ICP-MS, when PEO4 cells were treated with 0. 16 or 32 μM cisplatin for 5h following 24h pre-treatment with 20μM Vorinostat. Error bars show SEM. *** = p < 0.001, t.test. n=3. C) Cell survival of PEO1 and PEO4 cells on exposure to increasing concentrations of cisplatin for 24h, following pre-treatment with vehicle, or 20μM Vorinostat for 24h. Error bars show SEM. n=3.

## DISCUSSION

Cytotoxic, DNA targeting, chemotherapies, such as platinum based chemotherapy, have major impact on improving patient survival and are still the first line of treatment for many cancers. However, drug resistance is a major clinical problem leading to poor survival for patients. A diverse range of drug resistance mechanisms have been observed experimentally (39). However, with the exception of THBS1 and depending on the statistical parameters used, we observe relatively few consistent single gene changes in expression or chromatin conformation between the sensitive and resistant matched pairs. However, the ATAC-seq data of the HGSOC resistant lines in Figure 1 shows that the resistant lines clusters together rather than with their respective sensitive line, suggests there are common underlying epigenetic mechanisms involving changes in chromatin. The changes in chromatin conformation observed occur more frequently at intergenic regions rather than gene promoters and result in chromatin accessibility changes at COREs and hence change the super-enhancer epigenomic landscape.

Our data argue that epigenetic changes at super-enhancers during development of drug resistance is a common underlying mechanism that can drive expression changes in pathways such as those involved in Homologous Recombination repair leading to drug resistance. Such epigenetic changes may alter different resistance pathways and gene expression in a stochastic manner which can be selected on during tumour evolution. We observe enrichment for genes involved in the Fanconi anemia/BRCA DNA damage response pathway within the set of genes near super-enhancers gained during acquisition of drug resistance (36) and observe down regulation of their expression in the resistant PEO4 and PEA2 lines. This is consistent with loss of HR repair leading to platinum sensitivity.

Integrative genome-wide epigenomic and transcriptomic analyses of *in vitro* derived platinum-sensitive and –resistant ovarian lines identified key distal enhancers associated with platinum resistance and identified SOX9 as a critical super-enhancer regulated transcription factor that plays a critical role in chemoresistance in vitro of ovarian cancer cell lines (14). We also observe differences in super-enhancers in the in vitro derived A2780 and A2780/cp70 pair of lines, although these cluster separately from the HGSOC, in vivo, derived resistant lines (Figure 1). In our data, SOX9 is in the set of genes near COREs gained in the resistant PEO4 line, which is consistent with the data from Shang et al. showing increased H3K27ac around SOX9 in chemoresistant ovarian cancer cells (14).

THBS1 was the only gene consistently defined as downregulated in resistant lines using all methods of analysis of the RNAseq data.THBS1 is an important component of the extra-cellular matrix and has an important role in cancer development and regulating tumour cell behaviour (43). Although no evidence has been shown for THBS1 having a direct role in resistance to platinum drugs, it does have functions in tumour vasculaturisation, modulation of immune responses and is a pro-apoptotic factor, which may be important during tumour evolution during treatment. Further, ovarian cancer patients with high THBS1 expression and treated with platinum-based chemotherapy have longer survival (44) and is an independent prognostic factor in multivariate analysis.

In order to explore further the influence of chromatin organisation in cisplatin resistance we developed and applied Pt-exo-seq, a method for mapping cisplatin adducts genome-wide at base pair resolution. Genome-wide adduct formation distinguished the resistant PEO4 line from its isogenic sensitive counterpart, and was associated with the altered chromatin landscape we detected, highlighting altered adduct formation and distribution as a defining characteristic of these resistant ovarian cancer cells. Regions of the genome with fewer observed adducts in the resistant line were primarily around gene promoters, while regions with increased adducts were enriched for intergenic loci, and included super-enhancers which were reprogrammed in the resistant lines.

A limitation of the Pt-exo-seq study is that cells were treated for 5h to allow platinum crosslinks to form before DNA isolation for Pt-exo-seq and some DNA repair of the mono- or cross-linked adduct is likely to have occurred in this time. Transcription-coupled nucleotide excision repair may be responsible for the reduced persistence of adducts near gene promoters; a view supported by the difference in adduct formation at transcription start sites (TSS) of genes showing equally high expression in PEO1 and PEO4 (Supplementary Figure 9). Thus, variability in location of platinum in adducts detected in the genome may be due to differences in rate of DNA repair across the genome (22, 23), although may also be due to differences in chromatin accessibility for platinum damage. The current study has focussed on analysis of patient-derived platinum resistance, but the use of cell lines deficient in specific DNA repair pathways would help address some of these limitations.

It is well established that there is a difference in platinum DNA damage induction and repair in naked versus DNA bound to nucleosomes. For instance, studies on platinum damage at reconstituted chromatin show increased adduct formation predominantly in the nucleosome core. However, what the present study suggests is that it is not only the presence of nucleosomes that is important, but also whether they are in an open or closed conformation. Thus, regions of more compact chromatin had more adduct formation in resistant than in the sensitive HGSOC line, and this was particularly pronounced in intergenic regions. Furthermore, when examining COREs which were shared between PEO1 and PEO4, significantly more platinum adducts were detected in the resistant PEO4 line compared to COREs in the sensitive PEO1 line. Similarly, COREs which are lost in PEO4 incurred more adducts that the COREs which are maintained. This suggests that more platinum damage occurs at COREs in chromatin regions which become more nucleosome dense with the development of drug resistance, and that adducts at these loci are more tolerated by the cell, for instance due to increased error-prone translesion synthesis. While epigenetic approaches to overcome drug-resistance have been proposed and are undergoing clinical evaluation, they have generally failed to show clear patient benefit or in some cases have increased toxicity (40, 41). The use of non-specific epigenetic therapies that affect epigenetic marks throughout the genome will affect normal and tumour cells. This may explain the difficulties in improving therapeutic index in clinical trials of epigenetic therapies to overcome drug resistance. More targeted approaches based on reactivation of specific genes by CRISPR-epigenetic modulator fusions are now being explored (42) but are unlikely to overcome the diverse range of mechanisms involved in drug resistance if they only target specific genes. Targeting chromatin conformation in specific genomic contexts, rather than genome-wide, will be an important future strategy for epigenetic therapies to reverse drug resistance.

We have shown that chromatin changes, particularly at intergenic regulators of gene expression, are associated with *in vivo* derived drug resistance and Pt-adduct distribution in HGSOC. This has important implications for understanding the mechanisms of how tumour cells can adapt to treatment leading to drug resistance and treatment failure. These data suggest that resistance is driven by epigenomic changes which alter patterns of gene expression primarily through chromatin change at intergenic, super-enhancer regions rather than gene promoters. Therefore, new approaches are required that target the epigenome in terms of chromatin conformation in specific genomic contexts, rather than either the entire genome or single gene targeting.

## MATERIALS AND METHODS

### Cell lines and cell culture

The derivation of the cell lines has been previously described (24, 25). The identity of the lines was confirmed by STR profiling (Genetica). All cell lines were grown in RPMI-1640 (Sigma Aldrich) supplemented with glutamine (2 mM) (Gibco) and 10% fetal bovine serum (First Link). Cells were tested for mycoplasma monthly using the MycoAlertTM Mycoplasma Detection Kit (Lonza). Cell viability was assessed using MTS assay: CellTiter 96® AQueous One Solution Cell Proliferation Assay (Promega).

### ATAC-seq

Chromatin was extracted and digested as described (27). Libraries were sequenced on HiSeq 2500 (Illumina) using 100 bp paired-end reads. Quality assessment of libraries was performed using FastQC (http://www.bioinformatics.babraham.ac.uk/projects/fastqc). Duplicate reads were removed using the MarkDuplicates tool from the Picard tools suite (https://github.com/broadinstitute/picard). Reads were then aligned to the genome using Bowtie2 (28) with the parameters -X2000 and -m1. This ensured that fragments up to 2 kb were allowed to align (-X2000) and that only uniquely aligning reads were collected (-m1) as described (29). Fragment size data were collected using the CollectInsertSizeMetrics tool from the Picard suite. Reads were filtered against genomic blacklist regions, including mitochondrial sequences, defined by The ENCODE Project Consortium (30).

### RNA-seq

Libraries were prepared using the NEBNext Ultra Directional RNA Library Prep Kit for Illumina (NEB), according to the manufacturer’s instructions. Using the featurecounts function from the RSubread package in R (31), aligned reads were assigned to genes as counts, using a reference file of all known genes on the hg19 genome assembly obtained from the University of California Santa Cruz Genome Browser (32). Normalisation was carried out and differential gene expression was calculated using the limma package (33). RPKM values were calculated using the rpkm function from the limma package.

### Pt-exo-seq

Libraries for Pt-exo-seq were produced using a novel protocol for detecting platinum damage genome wide adapted from (20, 21). Following treatment with cisplatin (Hammersmith Hospital Pharmacy), DNA was isolated from cells using the Gentra PureGene kit (Qiagen). 1μg purified DNA was sheared using Bioruptor Pico (Diagenode) to sizes of ~ 150 bp. Fragment sizes were confirmed using TapeStation (Agilent). Fragmented DNA was then incubated at 30°C for 30 mins, with dNTPs, ATP, T4 DNA polymerase, Polynucleotide Kinase and DNA Polymerase 1 (Klenow fragment) to generate blunt ends before the reaction was stopped by incubation at 75°C for 20 mins. Fragments were then incubated with the annealed P7 adapters along with ATP, and the T4 DNA ligase. Adapter ligated fragments were purified and incubated with the phi29 DNA polymerase, along with dNTPs, to allow the overhang between the fragment end and the adapter to be filled in. Adapter ligated DNA was incubated with Lambda exonuclease for 1h at 37°C. RecJF and single stranded DNA binding protein (SSB) were added with Lambda exonuclease to maximise digestion efficiency. The platinum molecules which were still present in the fragment, adapter ligated, digested DNA were then displaced by incubation with sodium cyanide at 65°C for 2h. Complementary strand synthesis was carried out using adapters complementary to the P7 adapter sequence, along with the phi29 DNA polymerase and dNTPs. Finally, the P5 adapter was ligated to the purified, double stranded DNA by incubation for 60 min at 25°C and then 10 min at 65°C to inactivate the enzymes. Purified libraries were then amplified by PCR. The enzymes and oligonucleotides used for preparation of Pt-exo-seq libraries are listed in Supplementary Tables 1 and 2. Libraries were sequenced on HiSeq 2500 using 100bp single read sequencing. Library quality was assessed by FastQC before adapter trimming with BBDuK, and alignment to the hg19 genome assembly using bowtie2. Reads were filtered against the DAC Encode Blacklisted regions (30) and deduplicated as described for the ATAC-seq libraries. Aligned, deduplicated reads were assigned to genomic loci using a 1Kb sliding window, moved 900 bp along the genome. Coverage in these windows was then calculated for each platinum treated sample, against the mean coverage of three mock-treated replicates, as a log2 ratio. Further details on assay optimisation have been reported (19).

### Statistical analysis

#### ATAC-seq

Counts of ATAC-seq reads in genomic windows were calculated using the featurecounts function from the RSubread package (31). RPKM values were calculated using the rpkm function in the edgeR package (34). Reads were further filtered to remove PCR duplicates using the MarkDuplicates tool from the Picard Tools Suite (http://broadinstitute.github.io/picard/). The filtered, deduplicated reads were used in analyses which involved clustering the samples based on chromatin accessibility profiles.

ATAC-seq reads were assigned to 1Kb windows and these were filtered by requiring a depth >1 RPKM and < 1 SD above the median depth for all windows analysed. Coverage was normalised using the voom method to estimate the mean-variance relationship of the log of these counts. This generated a precision weight for each window which could be analysed using the limma Empirical Bayes analysis pipeline (33) in order to calculate log2FC and p values for changes in accessibility for each window. Windows were annotated according to their nearest genomic feature using HOMER (35). Enrichment odds ratios and confidence intervals were calculated for each class of genomic element in sets of windows showing significantly increased or decreased accessibility, compared to all analysed windows, using Fisher’s exact test.

Coverage around the TSS’ of genes was calculated in the same way as for the sliding window approach described above, except windows were 2 Kb centered around the TSS of genes of interest. Once FPKM values had been calculated, they were compared between cell lines using Students T-test.

ATAC-seq peaks were called in each ATAC-seq replicate using MACS2. These peaks were then used as input for the CREAM package (https://www.biorxiv.org/content/early/2017/11/21/222562), which calls clusters of cis-regulatory elements based on the genomic distribution of peaks. Genes within 100 Kb of called COREs were extracted from the Ensembl database using biomaRt. T statistics and odds ratios for differential expression of these genes were calculated from the RNA-seq data produced in this study.

#### Pt-exo-seq

Odds ratios for enrichment of platinum-target dinucleotides at the 5’ end of reads were calculated using Fisher’s exact test to compare platinum-target dinucleotide frequency at each position (−5 - +10 bp) in the read. Windows were filtered by intersection with the regions passing filters in the ATAC-seq data for the PEO pair (depth >1 RPKM and < 1 SD above the median depth for all windows analysed). Differentially damaged windows were defined as having a Log2 fold change in Pt-exo-seq signal:background ratio > ±2 and an FDR corrected p value < 0.05, based on a T-test between three replicates for each condition. Windows were annotated according to their nearest genomic feature using HOMER (35). Enrichment odds ratios and confidence intervals were calculated for each class of genomic element in sets of windows showing significantly increased or decreased accessibility, compared to all analysed windows, using Fisher’s exact test. Pt-exo-seq coverage around the TSS’ of genes was calculated in the same way as for the sliding window approach described above, except windows were 2 Kb centered around the TSS of genes of interest. Once RPKM values had been calculated, they were compared between cell lines using Students T-test.

#### MNase laddering following Vorinostat treatment

PEO4 cells were exposed to 20μM Vorinostat for 24h, nuclei isolated and incubated in micrococcal nuclease at a final concentration of 0.02 units/μl, in MNase buffer (New England Biolabs) at 37°C for 10 minutes before purification using the MinElute PCR cleanup kit (Qiagen). Isolated DNA was run on TapeStation High Sensitivity DNA tape (Agilent). The relative area of each peak was then used to calculate enrichment for mono-, di- and tri-nucleosome associated fragments. Differences in peak area were tested using students T-test.

### Quantification of DNA platination by Inductively Coupled Plasma Mass Spectrometry (ICP-MS)

Cells were treated with cisplatin for 5 h, before DNA was extracted. Platinum adduct levels were measured by ICP-MS using an Agilent 7900 mass spectrometer equipped with a micromist nebuliser and a cooled spray chamber (Agilent Technologies, USA). The assay was calibrated using platinum standards (Specure®, Alfa Aesar, Ward Hill, MA) and quality control samples with known platinum concentrations (ClinChek® Serum Controls, RECIPE, Munich, Germany). Platinum adducts were measured as previously described (26) in no-gas mode. Each measurement was made at 6 points on the mass peak in triplicate.

## Acknowledgements

This work is supported by an MRC doctoral training programme grant to JG, Cancer Research UK (A13086), Ovarian Cancer Action and Innovate UK. We would like to thank Prof Simon Reed, Cardiff University for advice on the platinum damage assays.

## Author contribution

JG contributed to data acquisition, methodology, analysis and writing (original draft), EL contributed to methodology, data acquisition and analysis, writing (review and editing), EC contributed to data curation, software and analysis, NM contributed to methodology and resources, LB contributed to funding acquisition and resources, IG contribuited to validation, RB contributed to conceptualisation, funding acquisition, resources, project administration, supervision and writing (original draft) JF and contributed to contributed to conceptualisation, supervision and writing (review and editing)

## Availability of data

The data discussed in this publication have been deposited in NCBI’s Gene Expression Omnibus (Edgar et al., 2002) and are accessible through GEO Series accession number GSE149147 (https://www.ncbi.nlm.nih.gov/geo/query/acc.cgi?acc=GSE149147).

## The paper explained

### Problem

Platinum-based chemotherapeutics, such as cisplatin and carboplatin, are clinically important first line therapies in the treatment of a wide variety of solid cancers. While many patients initially respond to platinum-based chemotherapy, they will eventually relapse with drug resistant disease that fails to respond to treatment leading to poor patient survival. Epigenetic mechanisms play a key role in the development of platinum-resistance. Understanding the chromatin and gene expression changes that occur in patient-derived platinum resistant tumours is essential for evaluating the importance of epigenetic change in tumour evolution and rational approaches to circumvent resistance.

### Results

We have investigated how genomic distribution of chromatin accessibilities alter during acquisition of resistance to carboplatin-based chemotherapy using matched ovarian cell lines from high grade serous ovarian cancer (HGSOC) patients before and after becoming clinically resistant to platinum-based chemotherapy. Resistant lines show altered chromatin accessibility at intergenic regions, but less so at gene promoters. Super-enhancers, as defined by clusters of cis-regulatory elements, at these intergenic regions show chromatin changes that are associated with altered expression of linked genes, with enrichment for genes involved in the Fanconi anemia/BRCA DNA damage response pathway. Further, genome-wide distribution of platinum adducts associates with the chromatin changes observed and distinguish sensitive from resistant lines. In the resistant line, we observe fewer adducts around gene promoters and more adducts at intergenic regions.

### Impact

The results have important implications for understanding the mechanisms of how tumour cells can adapt to treatment leading to drug resistance and treatment failure. These data suggest that resistance is driven by epigenomic changes which alter patterns of gene expression primarily through chromatin change at intergenic, super-enhancer regions rather than gene promoters. Therefore, new approaches are required that target the epigenome in terms of chromatin conformation in specific genomic contexts, rather than either the entire genome or single gene targeting.

